# Evaluation of DNA extracted from blood filter spots and eluates processed for enzyme linked immunosorbent assay (ELISA)

**DOI:** 10.1101/540633

**Authors:** Mark Andy Xatse, Jewelna Akorli, Irene Offei Owusu, Livingstone Gati, Michael David Wilson

## Abstract

Dried filter blood spots have become a significant blood collection method for screening individuals for clinical purposes. When used for ELISAs, they are normally discarded after the blood has been eluted. However, they may still be useful for extraction of DNA for molecular-based assays. The aim of this work was to determine the integrity of DNA extracted from filter paper spots from which blood has initially been eluted for ELISA with sample dilution buffer (SDB) and phosphate buffered saline (PBS). DNA was extracted from the eluted filter spots, the eluate, and dried blood filter spots (controls) using spin column extraction. The quality and quantity of the extracted DNA was assessed and used for PCR to further evaluate their usefulness in molecular assays. Concentration of DNA obtained was dependent on the buffer used for processing the filter blood blots. Accounting for the DNA concentration obtained from dried blood spots, which were used as controls, DNA extracted from the already eluted blood spots were 32 times higher in PBS than SDB processed filter paper. The ratio was even higher for the eluates, which were 57 times higher in PBS than SDS eluates. SDB eluates had significantly higher average DNA concentration than their eluted filter paper, but their purity ratios were similar. 85% PCR success rate was achieved with the DNA samples. Useful DNA can be extracted from blood spots after it has been eluted with SDB. Although the DNA concentration and purity may be low, the DNA could be useful for rather simple PCR assays.

**Author Summary:** Collection of blood onto filter paper has become an accepted method for screening individuals for clinical and public health purposes since the 1960s. This method of blood collection has become increasingly popular due to its ease and convenience in collection and transportation. The use of dried blood spots for clinical evaluations and research has become very significant. For research purposes, DBS when used for ELISAs are discarded after single use. DNA may however be extracted from the used filter blots and used for molecular assays. The concentration of DNA obtained may be low but simple assays like PCR could be done using the DNA extracted from the eluted filter spot.

## Introduction

The reliability and performance of molecular assays are strongly influenced by the quality and quantity of the starting template. The availability of high quality DNA from a large number of well characterized patients and healthy controls is a prerequisite for the success of genetic variation studies [1]. Conventionally, DNA used in clinical epidemiological studies is often obtained from peripheral blood samples [2,3]

Collection of blood onto filter paper has become an accepted method for screening individuals for clinical purposes. This type of specimen has been used for public health purposes since the 1960s [4] and has become increasingly popular due to its ease and convenience in collection and transportation. For example, to obtain blood samples from a baby, a few drops of blood from the baby’s heel are made to flow onto and fill a printed circle on a special filter paper. The blood dries under ambient atmospheric conditions, and the filter paper is mailed to a laboratory where a portion of the blood spot is punched out with a paper punch [4]. Biological markers that can be measured from whole blood, serum or plasma can be determined from dried blood spots [5]. This includes DNA, which is important for research or studies in genetics.

Blood spots have been used routinely since the 1960s[6] for neonatal screening, initially used for detecting phenylketonuria, and subsequently for other biochemical assays. They have also been used as a source of DNA for screening genetic abnormalities such as cystic fibrosis and haemoglobinopathies in newborns [7]. Filter blood spots are used for monitoring antibodies against several viral [8] and bacterial pathogens [9], storage of monoclonal antibodies [10] and, HIV screening [11]. Dried blood spots have been particularly useful for isolating parasite DNA in mapping the spread of drug resistance in malaria parasites [12].

The use of the parasite antigen Og4C3 ELISA is among several techniques used to identify infection with lymphatic filariasis (LF) in *Wuchereria bancrofti* endemic areas[13,14]. Typically, the Og4C3 ELISA is performed using serum or plasma, either immediately after the sample has been obtained, or more often, from frozen samples. However, the Og4C3 ELISA also offers an alternative application method through the use of dried blood spots (DBS) collected on filter paper; an inexpensive and convenient method that requires less space and less stringent refrigeration for transport and storage [15].

Filter spots used for Og4C3 ELISA assays are normally discarded after the blood has been eluted. However, in the advent of increased use of molecular assays for in-depth understanding of diseases, these eluted filter blots hold more information than what is only required for ELISAs. In some cases where molecular analyses are also required in a study besides ELISA, there may not be enough blood from an individual to have extra dried filter spots for DNA extraction. This study, therefore, aims to determine the integrity (quality and yield) of DNA obtained from filter paper spots from which blood has been eluted and the eluate intended for ELISA. We compared the quality and yield of DNA extracted with that from dried blood filter spots, and used the DNA samples in a PCR to ascertain the usefulness of the extracted DNA in a molecular assay

## Materials and Methods

### Sample collection and processing

A total of 50 TropBio filter disks (Cellabs, Australia) with blood spots collected from 50 subjects were retrieved from archives of a previous study at Noguchi Memorial Institute for Medical Research and randomly divided into two groups of 25. Two (2) ears of dried blood spots (DBS) were torn from each disk and an ear placed separately into 1.5 mL microcentrifuge tubes. The DBS were processed for DNA extraction as shown (Fig 1).

**Fig 1.**
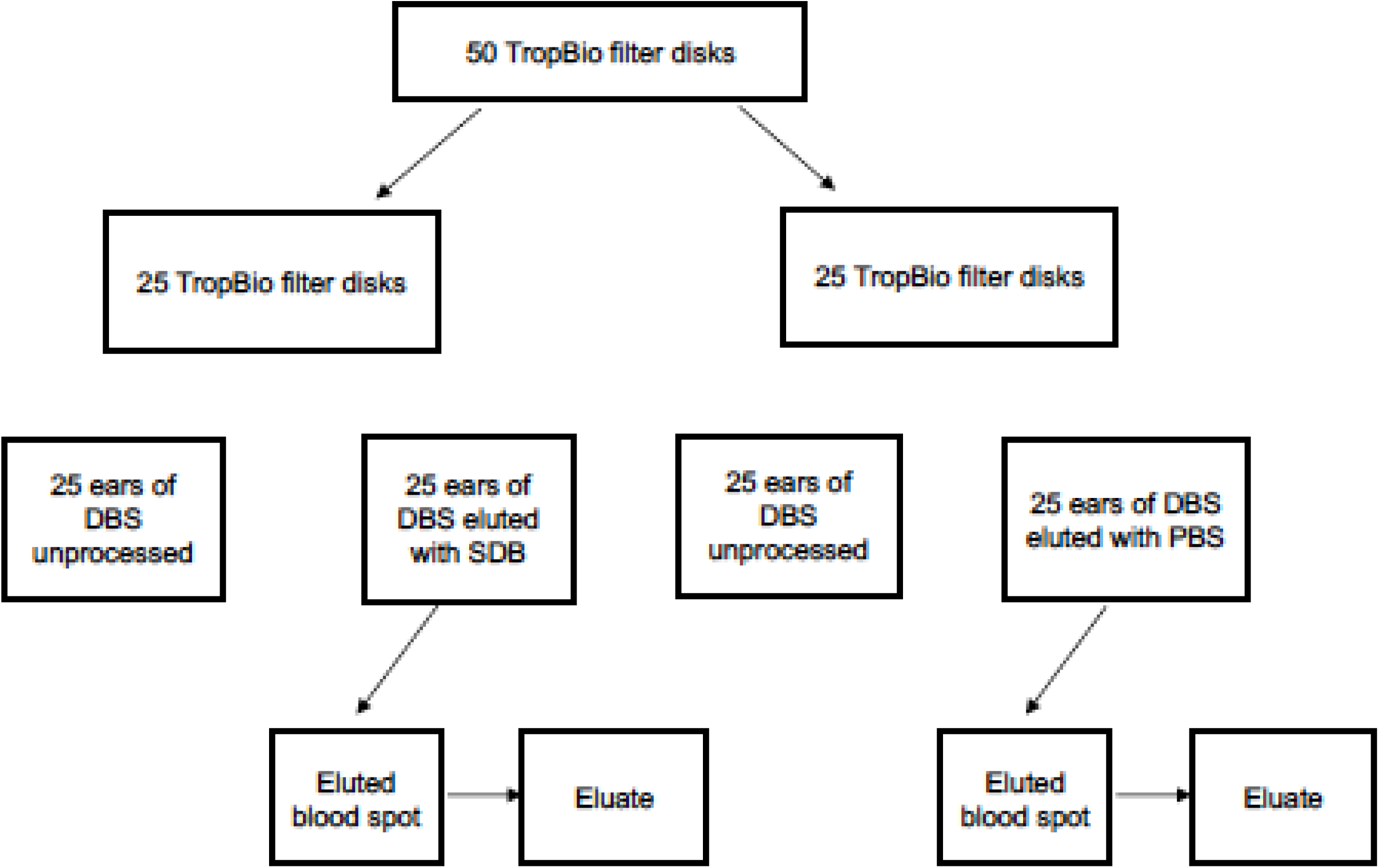
Sample processing and handling of filter blood blots prior to DNA extraction. DBS (dried blood spot); SDB (sample dilution buffer); PBS (phosphate buffered saline)

### Dry blood spot (DBS) elution and DNA extraction

250 µL of elution buffer (phosphate buffered saline or sample dilution buffer for Og4C3 ELISA) was added to one of the duplicate DBS in a 1.5mL microcentrifuge tube. The DBS was allowed to fully submerge in the buffer and then incubated overnight at 4°C on a gently rocking platform. The eluate was pipetted into a new 1.5 ml microcentrifuge tube, (leaving behind the eluted filter paper). DNA was extracted from the eluate, eluted filter paper and the dried unprocessed DBS using the Qiagen DNA Blood and Tissue kit according to the manufacturer’s instruction. DNA was eluted in 150 µL of buffer AE. DNA quantity and purity were measured using Qubit^®^ 2.0 fluorometer (Invitrogen, Life technologies) and NanoDrop 2000c Spectrophotometer (Labtech International Ltd, Thermo Scientific, UK). Samples were stored at −40°C for further analysis.

### Amplification of human ribosomal gene

PCR was performed to evaluate the quality of genomic DNA extracted. A 231bp region of the small subunit of human ribosomal gene (ssrDNA) was amplified [16,17] UNR-HUF, 5’-GAGCCGCCTGGATACCGC-3’ REV, 5’-GACGGTATCTGATCGTCTTC-3’ forward and reverse primers, respectively [18].

The PCR reaction consisted of 1X Phusion® High-Fidelity PCR Master Mix with HF Buffer, 4.0 mM MgCl_2_, 200 mM each of dNTPs, 0.0125 µM HUF primer, 0.075 µM REV primer and DNA template. The reaction was performed in a final volume of 12.5 µL using 2.5 µL DNA template. The reaction was performed in a thermal cycler (Applied Biosystems model 2720) with cycling conditions consisting of an initial denaturation phase of 98 °C for 1 min, 35 cycles at 98°C for 30 secs, 58°C for 1 min and 72°C for 1 min. The final cycle was followed by an extension time of 7 min at 72°C. The amplified fragment sizes were run on a 2% ethidium bromide stained agarose gel and viewed on a UV transilluminator (DS-30).

### Statistical analysis

Pairwise comparisons between treatments were performed using Kruskal-Wallis test with Benjamin, Krieger and Yekutieli correction for multiple comparisons. The significance level was set at 0.05.

### Ethics Statement

Ethical approval was obtained form the Ghana Health Service Ethical Review Committee for the samples collected for the main study. All sampled were anonymized before usage.

## Results

### Comparison of DNA concentrations between extraction templates

The aim of this work was to investigate the possibility of obtaining quality DNA from filter paper blots that have been previously been processed, and/or the eluate. Two different elution buffers (SDB and PBS) were used to obtain the eluate. Samples that were processed with SDB had an average DNA concentration of 0.15 ng/µl from the eluted blood spot and 0.08 ng/µl from the eluates (*p*= 0.01). Both concentrations were significantly lower than what was obtained from the dried blood spot (0.38 ng/µl) (eluted blood blot vs dried filter blot *p*= 0.03; eluate vs dried filter blot *p*< 0.0001) (Fig 2A). DNA samples extracted from filter blots eluted with PBS had an average concentration higher than the dried blood spot (control) samples (1.84 ng/µl vs 0.14 ng/µl) (*p*<0.0001). The DNA concentration from the eluates were on average similar to that from the eluted blood spots *(p*=0.10) (Fig 2B).

**Fig 2.**
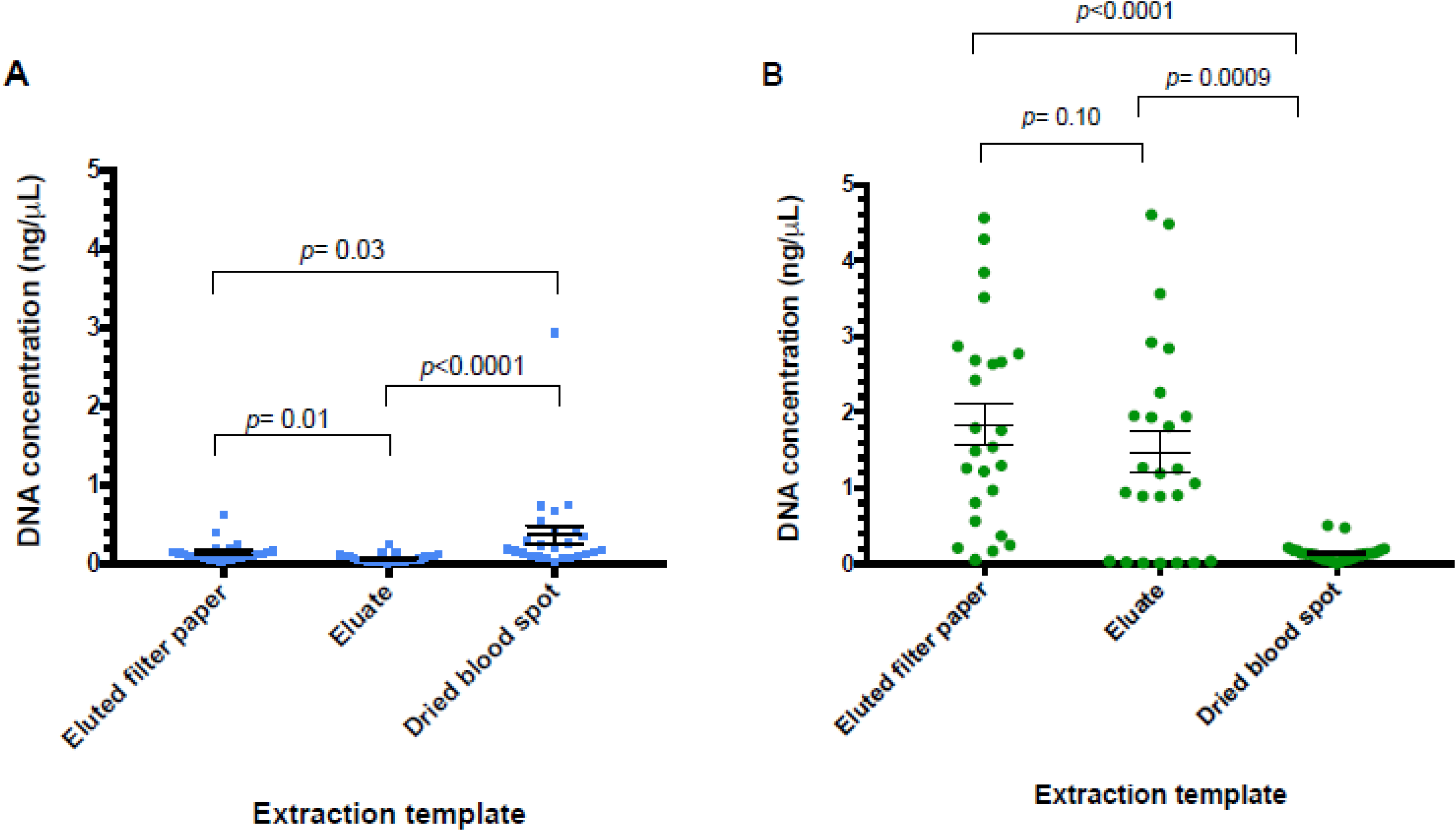
Comparison of DNA concentration extracted from eluted blood spot, dried blood spot and the eluate with sample dilution buffer (A) and phosphate buffered saline (B). The two elution buffers were compared to determine their influence on the resulting DNA concentration. Samples that had been processed with PBS prior to DNA extraction produced higher yields than SDB-treated samples (Fig 3). All SDB-processed samples were however similar to control dried filter blots from the PBS group (Fig 3).

**Fig 3.**
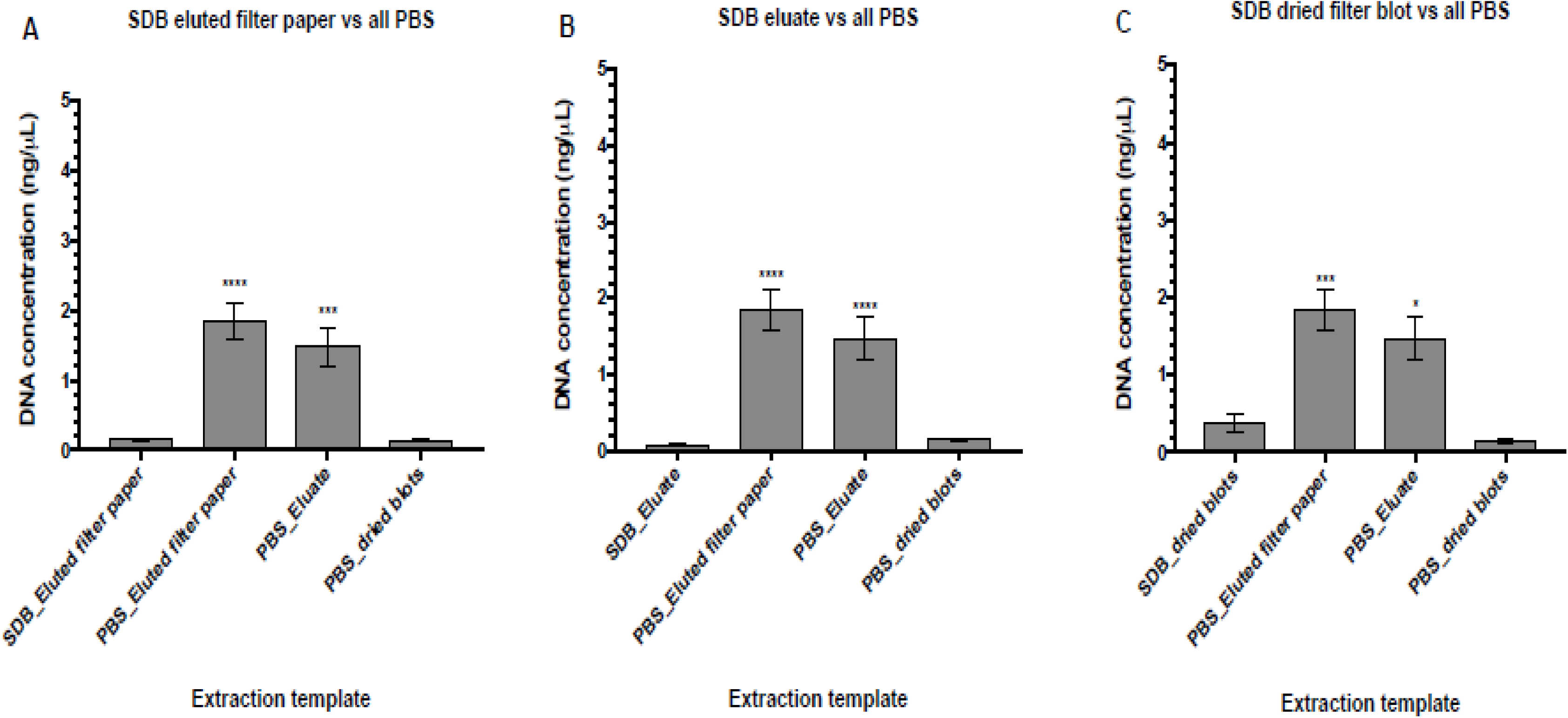
Comparisons of DNA yields between SDB and PBS processed samples. Only significant pairwise comparisons are shown with asterisks **** (<0.0001); *** (0.0001-0.001); ** (0.002-0.01); * (0.02-0.04).

### Purity ratios (A_260_/A_280_) for eluted blood spot, dried blood spot and the eluate

Purity was estimated using spectrophotometry as a measure of DNA usefulness in further molecular assays. An A_260_/A_280_ ratio between 1.7 and approximately to 2.0 is considered pure. None of the groups of samples extracted had an average purity ratio within the expected range (Fig 4). Altogether, 16% (24/150) of extracted DNA samples were estimated to be pure (Fig 4). The highest contribution of samples to this overall percentage was obtained from PBS-eluates (36%) and eluted blood spots (52%), and these measured higher in purity than any SDB-processed DNA sample in pairwise comparisons (Fig 4). Extractions from all dried blood spots were of low purity (Fig 4). Comparisons between the SDB-processed samples showed no difference in their average purity ratios (*p*<0.05) (Fig 4).

**Fig 4.**
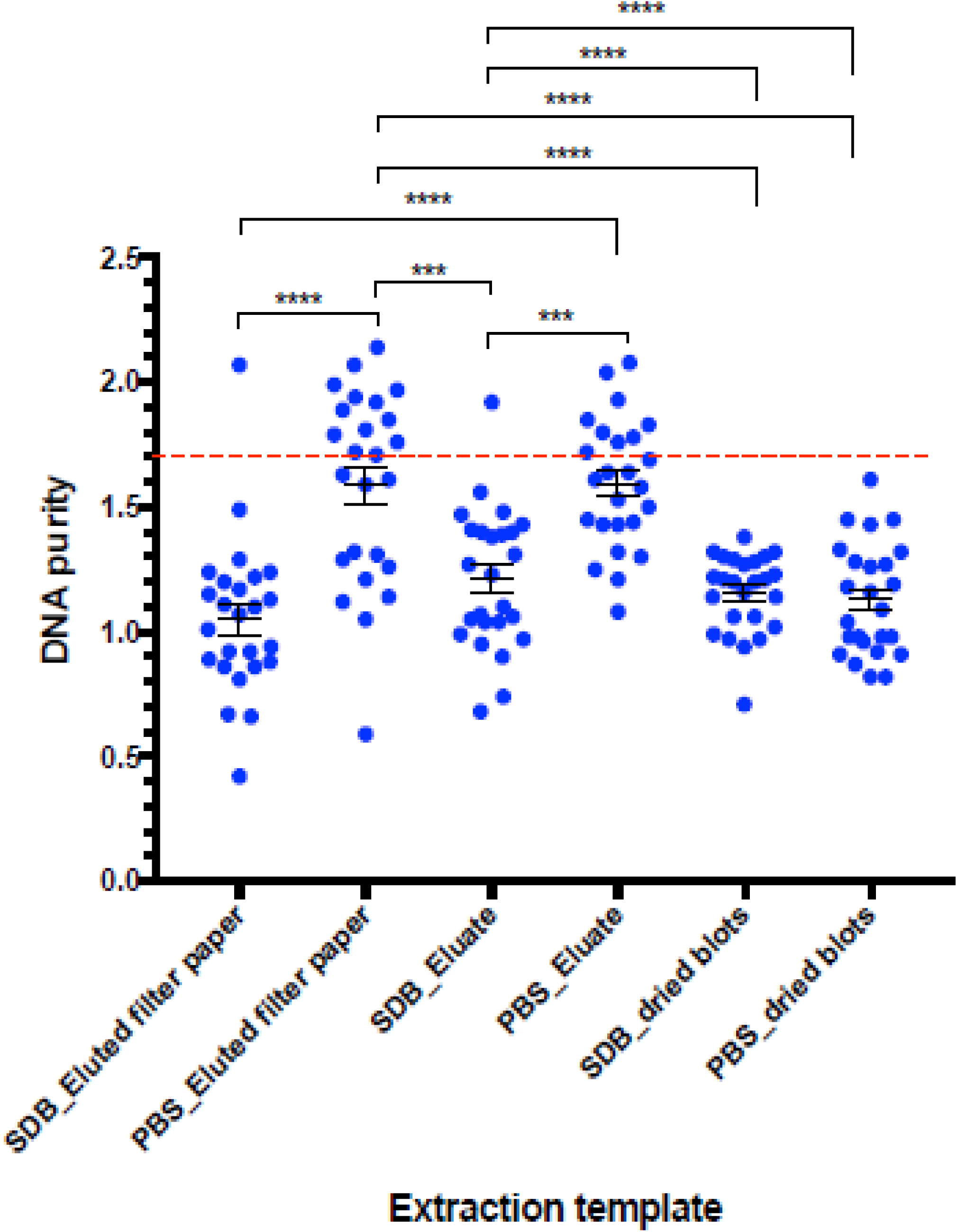
Purity ratios (A_260_/A_280_) of samples extracted from experimental templates. Error bars represent standard error of mean, and red dotted line indicates lower threshold for purity range (> 1.7). Only significant pairwise comparisons are shown with asterisks ****(<0.0001); ***(0.0001-0.001).

### PCR assay with extracted DNA sample

The human 18SrRNA could successfully be amplified in our samples (Fig 5), though DNA were of low concentrations and purity. Out of 150 DNA samples used in the assay, 127 produced positive PCR results (85%).

**Fig 5.**
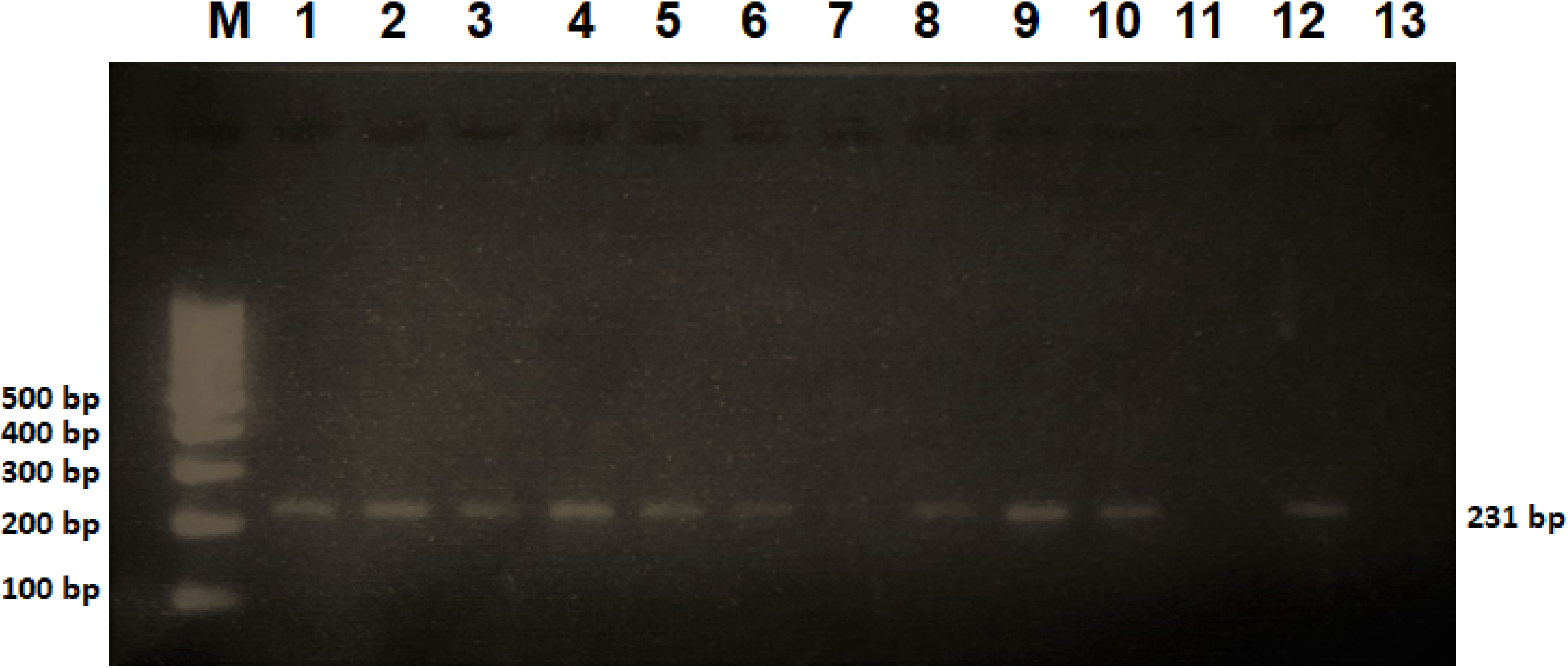
Gel electrogram of the amplified PCR products. Band size of 231 bp seen for samples in lanes 1 to 7 and 8 to 10. Lane 11 is a DNA from cultured *Plasmodium falciparum* 3D7 strain; Lane 12 is a positive control of DNA obtained from human blood; Lane 13 is a no template control (no DNA).

## Discussion

Obtaining blood samples from human subjects for research studies is expensive. Therefore, it is necessary to ensure that as much information required from samples can be extracted without having to return to the subjects for another blood collection. In the Lymphatic Filariasis Elimination Programme, blood samples are often collected on filter paper for ELISA-based assays such as Og4C3, Bm14 and Wb123. Although, the focus is not usually on DNA-based assays, this could later be an important inclusion in studying the molecular biology of parasites or infected humans. In this study, we considered the possibility of extracting useful DNA from filter blood papers that have already been processed for ELISA, and from the eluate which is commonly used for the ELISA assays. The DNA concentration and purity were compared among the starting materials to ascertain which gave better DNA integrity. Our results demonstrated that DNA concentration is dependent on the buffer used for processing the filter blood blots. Accounting for the DNA concentration obtained from dried blood spots, which were used as controls, DNA extracted from the already eluted blood spots were 32 times higher in PBS than SDB processed filter paper. The ratio was even higher for the eluates which were 57 times higher in PBS than SDS eluates.

The stability of double stranded DNA (dsDNA) can be affected by temperature, pH and ionic composition of solution (solvent)[19]. Salting-out has proven to be a cost-effective method for extracting DNA from whole blood, which gives good DNA yield for downstream analyses[20,21]. Phosphate buffered saline is a salt solution containing sodium chloride, sodium phosphate and potassium phosphate at a pH of 7.4. PBS does not only have high salt contents but it also has a pH which balances the salt concentration around cells, preventing osmosis [22]. On the contrary, the sample dilution buffer (SDB) consists of Tris buffer, sodium chloride, bovine serum albumin (BSA), and Tween buffer; it is of lower salt composition and has a pH of 8.0 [23]. Thus, PBS is expected to do better at salting-out DNA from the filter paper into solution (eluates) than SDB.

Besides DNA concentration, purity is critical in downstream applications such as PCR, and sequencing [24]. The purity of the DNA extracts further emphasized the preference on PBS over SDB-processed filter blots. Without the use of an appropriate solvent to put DNA into solution, most contaminants are retained on the filter paper which are later put into solution during the DNA extraction process. This is evident in the low purity of DNA from the dried filter spots (Figure 4). It is interesting to note that processed filter spots and their eluates were similar in purity.

## Conclusion

This study has established that DNA can be extracted from blood spots after it has been eluted with the sample dilution buffer used for ELISA-based assays. Although the DNA concentration (could be improved by reducing the elution buffer added at the end of the extraction process) and purity may be low, the DNA could be useful for simple PCR assays such as parasite DNA detection rather than sequencing which is highly sensitive to DNA concentration and purity.

## Acknowledgements

We acknowledge Nana Adjoa Pels and Frances McCarthy of the Department of Parasitology, UG-NMIMR for their assistance in the laboratory.

